# Intestinal microbiota of whitefish (*Coregonus* sp.) species pairs and their hybrids in natural and controlled environment

**DOI:** 10.1101/312231

**Authors:** Maelle Sevellec, Martin Laporte, Alex Bernatchez, Nicolas Derome, Louis Bernatchez

## Abstract

It is becoming increasingly clear that wild animals have never existed without symbiotic interactions with microbiota. Therefore, investigating relationships between microbiota and their host is essential towards a full understanding of how animal evolve and adapt to their environment. The Lake Whitefish (*Coregonus clupeaformis*) is a well-documented model for the study of ecological speciation, where the dwarf species (limnetic niche specialist) evolved independently and repeatedly from the normal species (benthic niche specialist). In this study, we compared the transient intestinal microbiota among five wild sympatric species pairs of whitefish as well as captive representatives of dwarf and normal species and their reciprocal hybrids reared in identical controlled conditions. We sequenced the 16s rRNA gene V3-V4 regions of the transient intestinal microbiota present in a total of 185 whitefish to (i) test for parallelism in the transient intestinal microbiota among sympatric pairs of whitefish, (ii) test for transient intestinal microbiota differences among dwarf, normal and both hybrids reared under identical conditions and (iii) compare intestinal microbiota between wild and captive whitefish. A significant effect of host species on microbiota taxonomic composition was observed in the wild when all lakes where analyzed together, and species effect was observed in three of the five species pairs. In captive whitefish, an influence of host (normal, dwarf and hybrids) was also detected on microbiota taxonomic composition and tens of genera specific to dwarf, normal or hybrids were highlighted. Hybrid microbiota was not intermediate; instead its composition fell outside of that observed in the parental forms and this was observed in both reciprocal hybrid crosses. Interestingly, six genera formed a bacterial core which was present in captive and wild whitefish, suggesting a horizontal microbiota transmission. Although diet appeared to be a major driving force for microbiota evolution, our results suggested a more complex interaction among the host, the microbiota and the environment leading to three distinct evolutionary paths of the intestinal microbiota.

## Introduction

Woese and Wheelis (1998) referred the Earth as a microbial planet, where macro-organisms are recent additions. Similar biochemical reactions between eukaryotes and prokaryotes in addition to the endosymbiotic theory suggest that eukaryotes evolved from prokaryotes [1, 2]. In addition, eukaryote-prokaryote cooperation fundamental to assemble more complex structures as multicellular organisms [3]. Hence, animals and plants have never been autonomous entities as they have always co-evolved in closed association with microbes [4]. Recent studies highlighted the substantial impact of microbiota on their host, for instance by causing the differential expression of hundreds of genes between germ-free and conventional organisms in several species [5, 6]. Furthermore, it is now clear that part of the microbiota is transmitted across generations in many animals and plants by various means [7]. In fishes in particular, the mother allocates antimicrobial compounds to the eggs before spawning [8, 9]. This maternal selection of bacteria influences the first bacteria that will be in contact with the sterile larvae during hatching [10]. Clearly then, a holistic understanding of macro-organisms biodiversity requires the investigation of their association with microbiota and their co-evolution.

The hologenome concept stipulates that the genome of the host and the microbiome (i.e. sum of the genetic information of the microbiota) act in consortium as a unique biological entity, that is, the holobiont [11]. Besides playing a role in their host adaptation to their environments [12], the microbiota may also be involved in reproductive isolation, either in the form of a pre-zygotic barrier by influencing the host’s mate choice by chemosensory signals [13, 14, 15] or post-zygotic barrier by producing genome and microbiome incompatibilities in hybrids, in accordance to the Bateson, Dobzhansky and Muller model of genetic incompatibilities [13, 16, 17]. Because the bacterial community of the gastrointestinal tract is implicated in many critical functions essential for development and immune responses, such as fermentation, synthesis and degradation functions, the intestinal microbiota could play an important role on its host’s adaptive potential [11, 12, 18].

Fishes as a group comprise the greatest taxonomic diversity of vertebrates and a major food resource for human populations [19, 20], yet little is known about the relationship with their microbiota and its evolution. A better understanding of fish intestinal microbiota is thus necessary towards filling the knowledge gap relative to the already well characterized mammals and insect microbiota [21]. The Lake Whitefish *(Coregonus clupeaformis)* is a well-studied system that represents a continuum in the early stage of speciation where sympatric species pairs of dwarf and normal species evolved independently in several lakes in northeastern North America [22, 23]. The normal species is specialized for using the trophic benthic niche, feeding on diverse prey as zoobenthos and molluscs. It is characterized by rapid growth, late sexual maturity and a long lifespan [24, 25]. In contrast, the dwarf whitefish is a limnetic specialist which feeds almost exclusively on zooplankton and is characterized by slower growth, early sexual maturation and shorter lifespan compared to the normal species. Previous transcriptomic studies revealed over-expression of genes implicated with survival functions (*e.g*. enhanced swimming performance for predator avoidance, detoxification) in dwarf whitefish, whereas normal whitefish show over-expression of genes associated with growth functions [22, 26]. Moreover, many other physiological, morphological and behavioral traits display parallel differences among these two whitefish species that correspond to their respective trophic specialization [22, 27-33]. Thus, the recent speciation and the clear trophic segregation make the whitefish species pairs an excellent model to study the role of intestinal microbiota in the context of ecological speciation. Two previous studies documented the variation of two microbial niches in Lake Whitefish species pairs, the kidney and the intestinal adherent communities [34, 35]. Although parallel pattern was highlighted between normal and dwarf species in the kidney communities, no clear evidence for parallelism was observed in the adherent intestinal microbiota. However, the water bacterial community was distinct from the adherent intestinal microbiota, indicating an intrinsic properties of the host microbiota [35].

Interestingly, there are emerging evidences that ingested environmental bacteria, including prey microbiota, play a significant role on the overall gut microbiota, either by stimulating colonization resistance or by providing additional functions to the host [36]. This allochthonous microbial community is called the transient microbiota. However, few studies tested parallelism patterns in fish intestinal microbiota [34, 37-41]. Also, the effect of the hybridization of two recently diverged species on microbiota composition is still poorly documented [42].

Therefore, the main goal of this study is to document the transient intestinal microbiota taxonomic composition of Lake Whitefish species pairs and their hybrids in natural and controlled environment. First, we investigated the transient intestinal microbiota in five wild species pairs of whitefish to estimate the within- and between-lake variation and tested for parallelism among transient intestinal microbiota. Secondly, we characterized the taxonomic composition of transient intestinal microbiota on dwarf, normal and first-generation hybrids reared in common garden in order to test the influence of the whitefish host on the microbiota in the same controlled conditions and under two different diets.

## Methods

### Sample collection for wild whitefish

Lake Whitefish were sampled in 2013 using gill nets in Cliff, Indian and Webster lakes in Maine, USA, and in East and Témiscouata lakes in Québec, Canada from May to July (Table 1). Fish were dissected in the field in sterile conditions as detailed previously [35]. Briefly, the ventral belly surface of fish was rinsed with ethanol, non-disposable tools were rinsed with ethanol and heated over a blow torch between samples. The intestine was cut at the hindgut end level (posterior part of the intestine) and the digesta were aseptically squeezed to collect the alimentary bolus. All samples of alimentary bolus were transported to the laboratory and kept at - 80°C until further processing.

**Table 1.** Number and locations of samples, sampling dates for each captive and wild whitefish populations or group. DD: dwarf whitefish, NN: normal whitefish, DH: hybrid F1 D♀×N♂, NH: hybrid F1 N♀×D♂.

| Origin | Form | Sample size | Sampling date | Coordinates |
| --- | --- | --- | --- | --- |
| Cliff | DD | 12 | June 13-14<sup>th</sup><br>2013 | 46°23'59"N, 69°15'11"W |
|  | NN | 12 |  |  |
| East | DD | 10 | July 2-4<sup>th</sup> 2013 | 47°11'15"N, 69°33'41"W |
|  | NN | 13 |  |  |
| Indian | DD | 12 | June 10-11<sup>th</sup><br>2013 | 46°15'32"N, 69°17'29"W |
|  | NN | 13 |  |  |
| Témiscouata | DD | 10 | May 28-30<sup>th</sup><br>2013 | 47°40'04"N, 68°49'03"W |
|  | NN | 14 |  |  |
| Webster | DD | 3 | June 12-13<sup>th</sup><br>2013 | 46°09'23"N, 69°04'52"W |
|  | NN | 12 |  |  |
| Common<br>Garden 1 | DD | 7 | November<br>12<sup>th</sup> 2013 to<br>June 09<sup>th</sup> 2014 | LARSA |
|  | NN | 5 |  |  |
|  | DH | 7 |  |  |
|  | NH | 6 |  |  |
| Common<br>Garden 2 | DD | 5 | November<br>12<sup>th</sup> 2013 to<br>June 10<sup>th</sup> 2014 | LARSA |
|  | NN | 4 |  |  |
|  | DH | 6 |  |  |
|  | NH | 6 |  |  |
| Common<br>Garden 3 | DD | 8 | November<br>12<sup>th</sup> 2013 to<br>June 11<sup>th</sup> 2014 | LARSA |
|  | NN | 6 |  |  |
|  | DH | 6 |  |  |
|  | NH | 8 |  |  |

### Experimental crosses, rearing conditions and sample collection for captive whitefish

In November 2013, 32 fish representing four cross types; dwarf (D♀ × D♂), Normal (N♀ × N♂), and their reciprocal hybrids (F1 D♀ × N♂ and F1 N♀ × D♂) were pooled together in three tanks (eight whitefish per form per tank). Experimental cross design was described previously [29, 31]. The protocol used for whitefish eggs fertilization and creating the parental generation is detailed in the Supplementary Data. Fish were separated in three tanks sharing the same experimental conditions (water, food, pH and temperature) for seven months. Juvenile whitefish were fed on two types of food; *Artemia* and dry food pellet BioBrood (Bio-Oregon^®^) [43, 44]. Although the fish were pooled without being marked, they were differentiated following genetic allocation to a host group based on mitochondrial (mtDNA) and nuclear DNA (Supplementary Data). In June 2014, fish were euthanatized with MS-222 and were dissected immediately in sterile conditions, with the same protocol and sterile tools as for wild whitefish. Samples were kept at 80°C until further processing. This study was approved under Institutional Animal Care and Use Committee protocol 2008–0106 at Laval University.

### Whitefish microbiota: DNA extraction, amplification and sequencing

The alimentary boluses from captive and wild fish were extracted using a modification of the QIAmp© Fast DNA stool mini kit (QIAGEN). To maximize DNA extraction of gram-positive bacteria, temperature and time were increased during the incubation steps and all products used were doubled (Proteinase K, Buffer AL and ethanol 100%). Thus, 1200 μl were transferred into the column (in two subsequent steps) and bacterial DNA was eluted from the column with 100 μl of ultrapure water (DEPC-treated Water Ambion^®^). DNA extractions were quantified with a Nanodrop (Thermo Scientific) and stored at –20°C until use. Five blank extractions were also done as negative controls.

In order to construct the community library, a region ~250 bp in the 16S rRNA gene, covering the V3–V4 region, was amplified using specific primers with Illumina barcoded adapters Bakt_341F-long and Bakt_805R-long in a dual indexed PCR approach [45]. The PCR amplification comprised 50 μl PCR amplification mix containing 25 μl of NEBNext Q5 Hot Start Hifi PCR Master Mix, 1 μl (0.2 μm) of each specific primers (Bakt_341F-long and Bakt_805R-long), 15 μl of sterile nuclease-free water and 8μl of specify amount of DNA. The PCR program consisted of an initial denaturation step at 98°C for 30s, followed by 30 cycles, where one cycle consisted of 98°C for 10 s (denaturation), 56°C for 30 s (annealing) and 72°C for 45s (extension), and a final extension of 72°C for 5 min. Negative and positive controls were also performed using the same program.

All PCR results, including negative controls, were purified using the AMPure bead calibration method. The purified samples were quantified using a fluorometric kit (QuantIT PicoGreen; Invitrogen), pooled in equimolar amounts, and sequenced paired-end using Illumina MiSeq at the Plate-forme d’analyses génomiques (IBIS, Université Laval, Québec, Canada). To prevent focusing, template building, and phasing problems due to the sequencing of low diversity libraries such as 16S amplicons, 50% PhiX genome was spiked in the pooled library.

### Amplicon analysis

Raw forward and reverse reads were quality trimmed, assembled into contigs for each sample, and classified using Mothur v.1.36.0 following the protocol of MiSeq SOP (https://www.mothur.org/wiki/MiSeq_SOP) [46, 47]. In brief, contigs were quality trimmed with several criteria. First, a maximum of two mismatches were allowed when aligning paired ends and ambiguous bases were excluded. Second, homopolymers of more than eight, sequences with lengths less than 400 bp and greater than 450 bp, sequences from chloroplasts, mitochondria and nonbacterial were removed. Thirdly, chimeric sequences were found and removed using the UCHIME algorithm [48]. Moreover, the database SILVA was used for the alignment and the database RDP (v9) was used to classify the sequences with a 0.03 cut-off level. The Good’s coverage index which was used to evaluate the quality of the sequencing depth, was estimated in Mothur [49].

### Statistical analyses

The analyses for the captive and wild whitefish microbiota were performed with Mothur and Rstudio v3.3.1 [50]. We first constructed a matrix of taxonomic composition (wild and captive included) with the number of Operational taxonomic units (OTUs) after merging them by genus. The bacterial genera were considered as variables and fish as objects according to Mothur taxonomy files (stability.an.shared and stability.an.cons.taxonomy).

We then investigated the microbiota difference between the captive and wild whitefish using a network analysis. A Spearman’s correlation matrix following a Hellinger transformation on the matrix of taxonomic composition was performed to document interactions between all captive and wild whitefish microbiota. More precisely, a Spearman’s correlation value (threshold ≥0.5), a P-value and Bonferroni correction was calculated for each sample. The network was visualized using Cytoscape v3.2.1 [51], where nodes were illustrated in two different versions: (i) according to their sampling sites (eight groups: five lakes and three tanks) and (ii) according to their genetic group (the two wild species pairs and the four captive groups: dwarf, normal, reciprocal hybrid F1 D♀N♂, and hybrid F1 N♀D♂). We also tested for the effect of captivity (wild and captive conditions) on whitefish microbiota taxonomic composition (PERMANOVAs; 10,000 permutations) and alpha diversity (inverse Simpson diversity) with an ANOVA following a fitted Gaussian family generalized model (GLM) [52]. This was performed on all fish, on dwarf whitefish only and on normal whitefish only. Finally, we documented the bacterial core of whitefish by identifying the bacterial genera present in 80% of all fish.

To document variation within and among wild whitefish populations, we tested for an effect of ‘host species’, ‘lake’ and their interaction, with ‘body mass’ as a covariate on the taxonomic composition, using a permutational analysis of variance (PERMANOVA; 10,000 permutations). This procedure was run for each of the five lakes independently after removing the explanatory variable ‘lake’ of the analysis. The ‘host species’, ‘lake’ effects and their interaction on the inverse Simpson diversity were also tested using an analysis of variance (ANOVA) following a fitted Gaussian family generalized model (GLM). Allometric effect on inverse Simpson diversity was first tested with a linear regression on body mass. As for the taxonomic composition, we ran this procedure for each lake independently. We also tested for differences in microbiota taxonomic composition between dwarf and normal wild whitefish by means of a discriminant analysis. First, a principal component analysis (PCA) on the transformed Hellinger matrix was conducted to avoid collinearity and over fitting problem. Only the axes explaining at least 1% of the variation were kept for the discriminant analysis, which was validated according to the method described in Evin *et al.* [53]. Furthermore, principal coordinates analyses (PCoAs) was built on a Bray-Curtis distance matrix after a Hellinger transformation to visualize variation at the genus level between dwarf and normal wild whitefish among and within the lakes [54, 55].

We then tested for differences in taxonomic composition between the four captive groups by investigating the effect of ‘host group’ (Dwarf, Normal, hybrid F1 D♀N♂, and hybrid F1 N♀D♂), ‘diet’ and their interaction with ‘body mass’ and ‘tank’ as covariates (PERMANOVA; 10,000 permutations). The effect of diet was added in the analysis because fish bolus exhibited a clear distinction between two observed feeding habits during the controlled experiment (A: feeding on a mix of dry food and *Artemia,* B: feeding on *Artemia* only). For the alpha diversity, the effect of ‘host group’, ‘diet’ and their interaction on the inverse Simpson diversity were tested with a mixed effects linear random model using the ‘nlme’ package in R, with tank as a random effect and individual fish nested within tank [56]. As for the analyses on wild whitefish, we first tested for an allometric relationship with body mass using a linear regression and used the residuals in all cases showing a significant relationship. Principal coordinates analyses (PCoAs) built on a Bray-Curtis distance matrix after a Hellinger transformation were also used to visualize variation at the genus level as described above. Discriminant analyses were also performed on captive whitefish but results were not displayed because of a negative cross-validation according to Evin *et al.* [53]. Moreover, in order to test for the presence of bacterial genera that were private to any of the captive whitefish group, we used the Metastats software with standard parameters (p ≤ 0.05 and number of permutations = 1000) to detect differential abundance of bacteria at the genus level between two host populations [57]. Four Metastats analyses were performed on the captive whitefish between: Dwarf vs. Normal, Dwarf vs. hybrid F1 D♀N♂, Normal vs. hybrid F1 N♀D♂, and hybrid F1 D♀N♂ vs. hybrid F1 N♀D♂.

## Results

### Sequencing quality

A total of 2,498,271 sequences were obtained after trimming for the entire data set composed of 185 whitefish intestinal microbiota (67 dwarf whitefish, 79 normal whitefish and 39 hybrids whitefish) from wild and captive populations (Table S1). A total of 189,683 operational taxonomic units (OTUs) were identified with a 97% identity threshold, representing 710 bacterial genera. Two wild whitefish samples were removed because of low coverage and low number of sequences and 22 captive whitefish samples were not included from various reasons (empty intestine, no PCR amplification or ambiguous genetic allocation to a host group).

The average Good’s coverage estimation for all intestinal microbiota (wild and captive whitefish) was 92.3±7.6%. While this may seem relatively low, the Good’s coverage from wild whitefish microbiota (n=111) and captive whitefish microbiota with a diet of *Artemia* only (n=27) were respectively 95.4±2.8% and 98.2±1.4%, thus indicating a good sequencing quality of our data. The low Good’s coverage essentially came from captive whitefish microbiota with a mixed diet of *Artemia* and dry food (n=47), with a coverage index of 82.8±3.4%. This data was considered reliable for further analysis for three reasons. First, this second diet group was composed of 341 bacterial genera in which the distribution showed an unusual high abundance (i.e. number of reads) for a few genera (Table S2), which is known to decrease the Good’s coverage which is defined as 1-(Number of OTUs that have been sampled once / total number of sequences) [49]. Second, the Illumina MiSeq sequencing was performed in the same run for all samples, thus supporting the absence of sequencing problem given the excellent coverage obtained for the other groups. Third, a low Good’s coverage is supposed to reflect a low number of sequences per sample because of the different filtration steps which eliminated reads generated by poor quality sequencing. Here, the low Good’s coverage observed in the captive group that fed on a mixed diet shows a total number of sequences per sample similar to the other captive group (Table S2).

### Wild *versus* captive whitefish intestinal microbiota

We observed a pronounced differentiation in intestinal microbiota between wild and captive whitefish based on the network analysis among all samples (Fig. 1). More specifically, a first group comprised of the wild whitefish (only one dwarf and two normal all from East Lake were excluded from this group). There was no clear pattern of differentiation between wild dwarf and normal whitefish microbiota (Fig. S1) but all wild populations tended to cluster together and distinctively from captive fish. The second and third groups were composed by all captive whitefish with few interactions observed between them despite the fact that they both comprised fish from all four groups (dwarf, normal and both reciprocal hybrids). This second level of differentiation was based on diet variation between the two groups of captive fish (Fig. 1). The differentiation between the wild and the captive fish was also supported by a significant effect of captivity on taxonomic composition (PERMANOVA, *P* < 0.001; Table 2) when performing analysis using all fish, dwarf only and normal only, as well as on alpha diversity when using all fish (ANOVA, *P* < 0.001; Table S3). Furthermore, although the relative abundance of major phyla of *Firmicutes, Proteobacteria, Actinobacteria* and *Planctomycetes* was similar between wild and captive whitefish, the bacterial abundance was clearly different between them (Fig. 2). Finally, among the 710 bacterial genera found among all captive and wild whitefish, six bacterial genera were shared by all fish: *Acinetobacter*, *Aeromonas*, *Clostridium*, *Legionella*, *Methylobacterium* and *Propionibacterium,* which together constitute the core intestinal microbiota defined as the microbial component shared by 80% of the samples.

**Figure 1.**
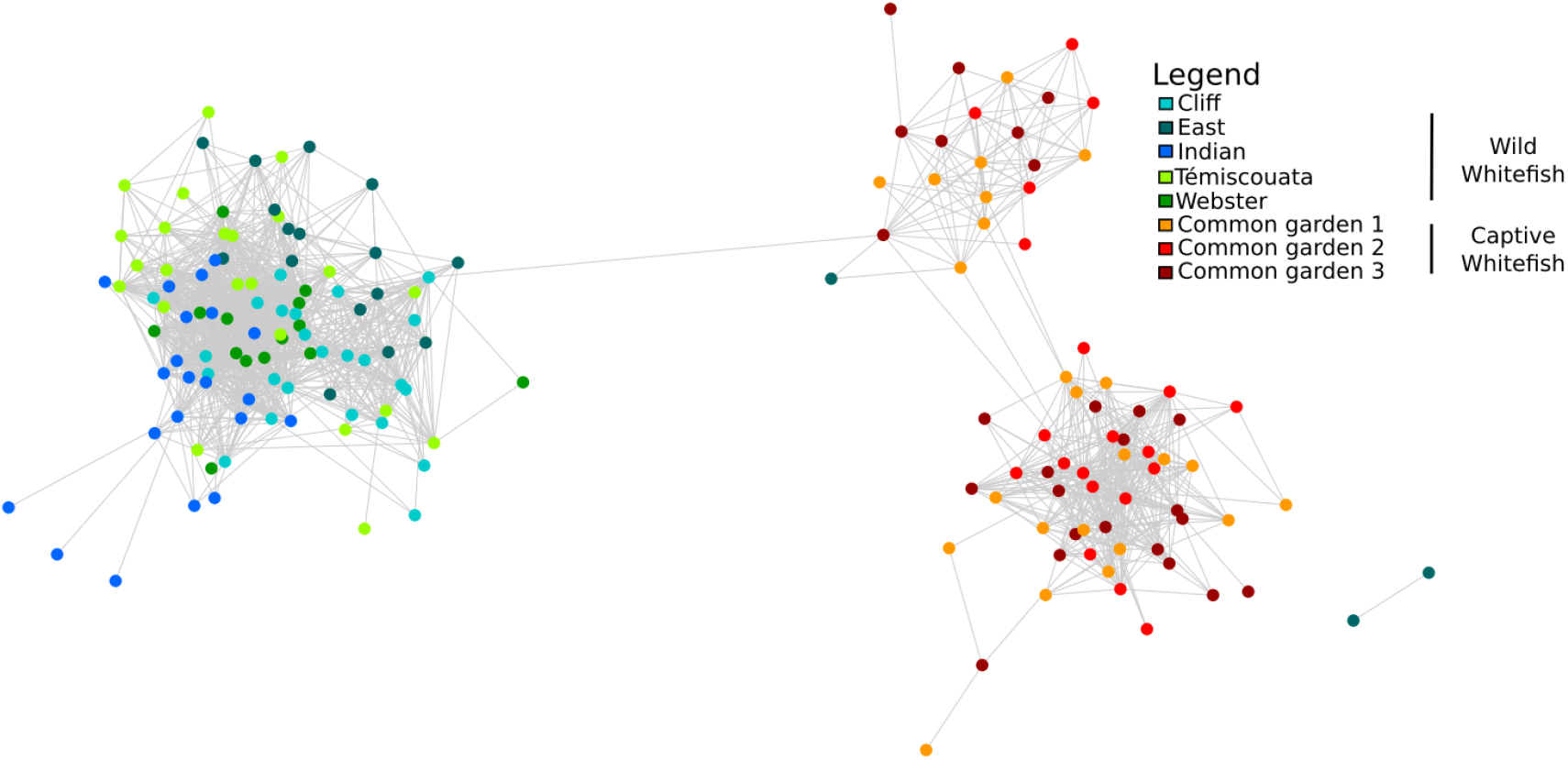
Network analysis of intestinal microbiota of dwarf and normal wild whitefish and intestinal microbiota of dwarf, normal and hybrids captive whitefish. Each node represents either a dwarf, normal or hybrid whitefish microbiota. The connecting lines between two samples represent their Spearman index correlation.

**Table 2.** Summary of PERMANOVA test statistics on microbiota taxonomic composition. First, the fish group “wild” refers to the analysis of effect of host species (dwarf and normal), lake (Cliff, East, Indian, Témiscouata and Webster) and its interaction with body mass as a covariate on all wild fish. Second, the fish group “All lakes” tests the host species and body mass as a covariate is treated for each lake separately. Third, the fish group “captive” refers to the analysis of effect of host group (dwarf, normal, hybrids F1 D♀N♂ and F1 N♀D♂), diet *(Artemia* only and mixed diet of live *Artemia* with dry food) and its interaction with body mass and tank as covariates on all captive fish. Fourth, the fish group ‘both’ refers to the effect of captivity (wild and captive) and body mass as covariate on all fish, dwarf only and normal only. F-Value: value of the F statistic, R^2^: R-Squared Statistic, Pr(>F): p-value.

| Fish group | Source of variation | PERMANOVA |  |  |
| --- | --- | --- | --- | --- |
|  |  | F-Value | R<sup>2</sup> | Pr(>F) |
| WILD |  |  |  |  |
| All lakes | Species | 2.350 | 0.017 | 0.006 |
|  | Lake | 6.744 | 0.197 | <0.001 |
|  | Species:Lake | 1.927 | 0.056 | <0.001 |
|  | Body mass | 1.628 | 0.012 | 0.067 |
| Cliff Lake | Species | 5.253 | 0.180 | <0.001 |
|  | Body mass | 2.914 | 0.100 | <0.001 |
| East Lake | Species | 1.889 | 0.085 | 0.047 |
|  | Body mass | 1.165 | 0.053 | 0.291 |
| Indian Lake | Species | 2.032 | 0.083 | 0.041 |
|  | Body mass | 1.582 | 0.064 | 0.105 |
| Témiscouata Lake | Species | 0.741 | 0.033 | 0.732 |
|  | Body mass | 0.920 | 0.041 | 0.447 |
| Webster Lake | Species | 0.858 | 0.057 | 0.562 |
|  | Body mass | 2.142 | 0.143 | 0.015 |
| CAPTIVE |  |  |  |  |
|  | Group | 1.985 | 0.043 | 0.033 |
|  | Diet | 58.955 | 0.427 | <0.001 |
|  | Species:Diet | 1.557 | 0.034 | 0.108 |
|  | Body mass | 1.990 | 0.014 | 0.084 |
|  | Tank | 1.649 | 0.024 | 0.102 |
| BOTH |  |  |  |  |
| All fish groups | Captivity | 64.457 | 0.260 | <0.001 |
|  | Body mass | 3.481 | 0.014 | 0.001 |
| Dwarf | Captivity | 28.245 | 0.289 | <0.001 |
|  | Body mass | 4.517 | 0.046 | <0.001 |
| Normal | Captivity | 16.371 | 0.180 | <0.001 |
|  | Body mass | 1.917 | 0.021 | 0.035 |

**Figure 2.**
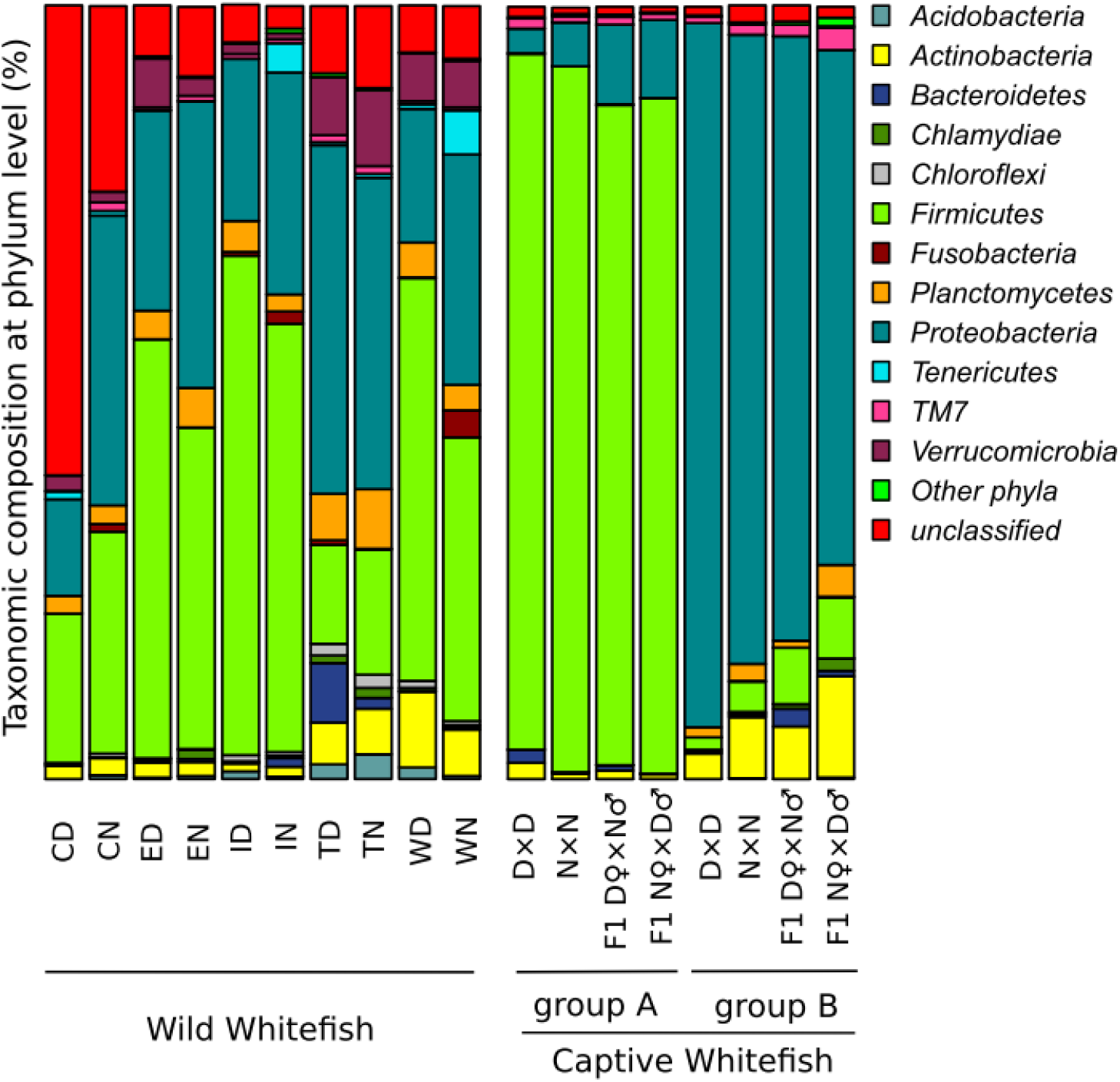
Relative abundance of phyla representatives found in intestinal microbiota for dwarf and normal wild whitefish in each lake, as well as in intestinal microbiota for dwarf, normal and hybrids whitefish in controlled condition. Taxonomy was constructed with the database Silva and MOTHUR with confidence threshold at 97%. For the wild whitefish, lakes are represented as C: Cliff, E: East, I: Indian, T: Témiscouata, W: Webster and the whitefish species is represented as D: dwarf and N: normal. For the captive fish, normal whitefish, dwarf whitefish and hybrids are represented as N×N, D×D, F1 D♀×N♂ and F1 N♀×D♂ respectively. Diet group A (*Artemia* + dry food) and B (*Artemia*).

### Wild dwarf and normal whitefish microbiota

At the phylum level, dwarf and normal wild whitefish transient intestinal microbiota was characterized by identical dominant phyla with a similar bacterial abundance (Fig. 2). The five main phyla for intestinal whitefish species microbiota were *Firmicutes, Proteobacteria, Planctomycetes, Verrucomicrobia* and *Actinobacteria.* However, taxonomic variation between dwarf and normal species were observed for less dominant phyla. For example, *Tenericutes* and *Fusobacteria* were more represented in normal whereas *Bacteroidetes* was more represented in dwarf whitefish. We observed a more pronounced influence of the lake whereby dwarf or normal microbiota within a given lake shared more similarities than microbiota from different lake populations within a same species (Fig. 2).

Although no effect of lake or species on alpha diversity was observed (Table S3), there was a significant effect of both lake and host species on taxonomic composition (Table 2). The discriminant analysis performed on all wild whitefish also confirmed this overall difference between dwarf and normal intestinal microbiota albeit with overlap between them (Fig. 3). Within each lake, the PERMANOVA tests revealed significant differences between dwarf and normal whitefish in three lakes (Cliff, East and Indian lakes) whereas no significant difference was observed in Témiscouata and Webster lakes (Table 2). Together, these observations suggested that the lake effect was more important than that of the host species. This is also supported by the PCoA analyses that revealed no global differentiation between all dwarf and normal whitefish intestinal microbiota (Fig. 4-A). Host effect is nevertheless also supported in lake specific PCoAs based on partially overlapping 95% confidence interval in Cliff, East and Indian lakes (Fig. 4-B:D). Still, complete overlap was observed in Témiscouata lake (Fig. 4-E) and results are ambiguous in Webster lake considering the low sample size (Fig. 4-F).

**Figure 3.**
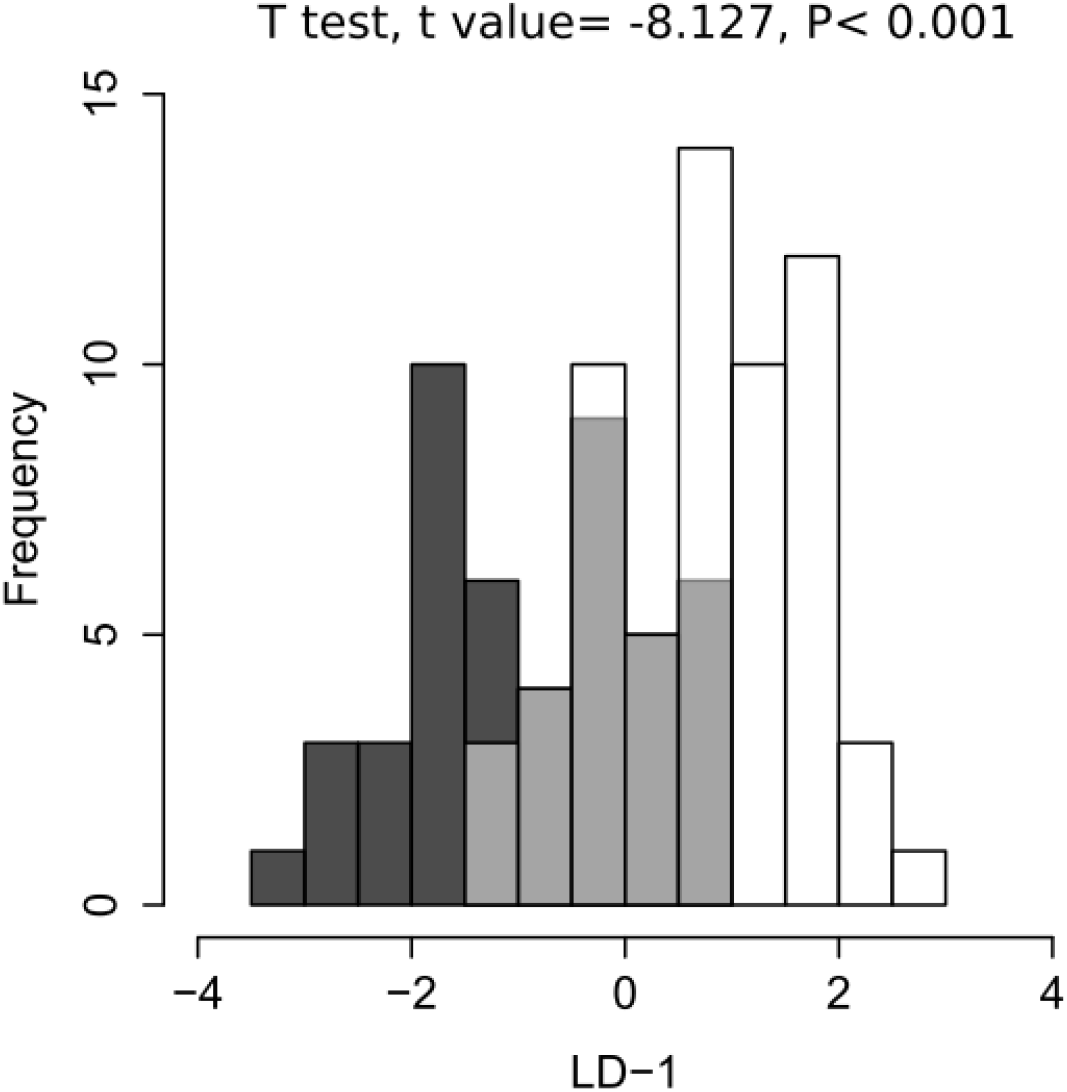
Discriminant analysis histogram of all wild whitefish microbiota. This discriminant analysis was performed on the axes of principal component analysis (PCA) and T-tests were performed on the results of the discriminant analysis. Dwarf and normal whitefish are represented by the black and white bars, respectively. Dwarf and normal whitefish with overlapping discriminant scores are shown in grey.

**Figure 4.**
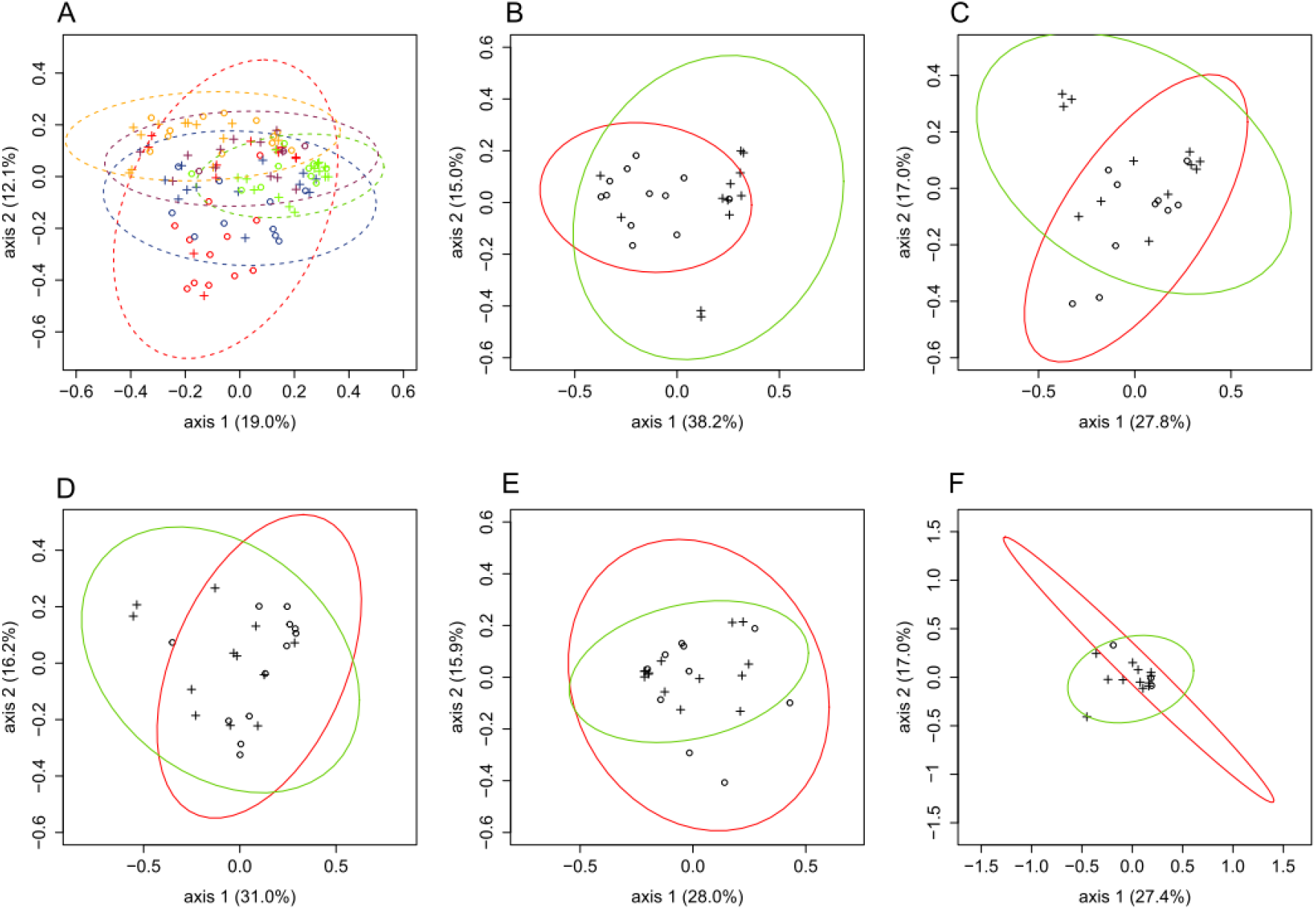
Principal coordinate analyses (PCoAs) within- and between lakes for the wild whitefish microbiota. These PCoAs are based on Jaccard index after a Hellinger transformation. Ellipses of 95% confidence are illustrated and were done with dataEllips using R car package. A: comparison among all wild whitefish populations from the five lakes. Each lake analyzed is represented by a different symbol and ellipse color: Cliff Lake (red), East Lake (blue), Indian Lake (orange), Témiscouata Lake (green) and Webster Lake (purple) and whitefish species is represented by symbols: Dwarf (circle) and Normal (cross). B-F: comparison between Dwarf and Normal whitefish microbiota within each lake. Cliff Lake, East Lake, Indian Lake, Témiscouata Lake and Webster Lake are represented by B, C, D, E and F respectively. Whitefish species is represented by different symbols: dwarf (circle) and normal (cross); ellipses of 95% confidence are illustrated and were done with dataEllips using R car package. the red and green ellipses represent the dwarf and normal species, respectively.

### Pure and hybrid whitefish microbiota in controlled environment

Although all fish were exposed to the same environment and the same food (both *Artemia* and dry fish food), we observed that some whitefish did not feed on the dry fish food and ate only live *Artemia*. As for the network analysis, the two distinct diet groups were evidenced by a significant effect of diet on both taxonomic composition microbiotas (PERMANOVA, P < 0.001; Table 2) and alpha diversity (ANOVA, *P* = 0.001; Table S3). The PCoA analysis clearly separated two distinct clusters on axis one corresponding to the two diet groups and independent of the genetic background (either pure forms or hybrids) (Fig. 5). Furthermore, the mixed diet group was dominated by *Firmicutes* and the *Artemia* diet group was dominated by *Proteobacteria* (Fig. 2). The five main phyla composing the microbiota of the mixed diet group were in ranked order *Firmicutes, Proteobacteria, Actinobacteria, Bacteroidetes* and *TM7* whereas the five main phyla of the *Artemia* diet group were *Proteobacteria, Actinobacteria, Firmicutes, Planctomycetes* and *TM7.* Within the mixed diet group, lower abundance for Firmicutes, but higher for *Proteobacteria* was observed in reciprocal hybrids in comparison to dwarf and normal whitefish. Interestingly, the opposite pattern was observed for the *Artemia* diet group (i.e. hybrids bacterial abundance was higher for *Firmicutes* but lower for *Proteobacteria*). The PERMANOVA tests also revealed a significant effect of host group (Table 2). The PCoA analysis within each of the two diet groups highlighted a modest differentiation based on overlapping 95% confidence interval between hybrids and pure whitefish (Fig. 5). In the mixed diet group, dwarf and normal ellipses were mostly aligned on the second axis whereas the ellipses of the two hybrid groups were mostly aligned on the first axis. The inverse pattern was observed in the *Artemia* diet group with the ellipses of the pure whitefish those of hybrid whitefish aligned on the first and second axis, respectively.

**Figure 5.**
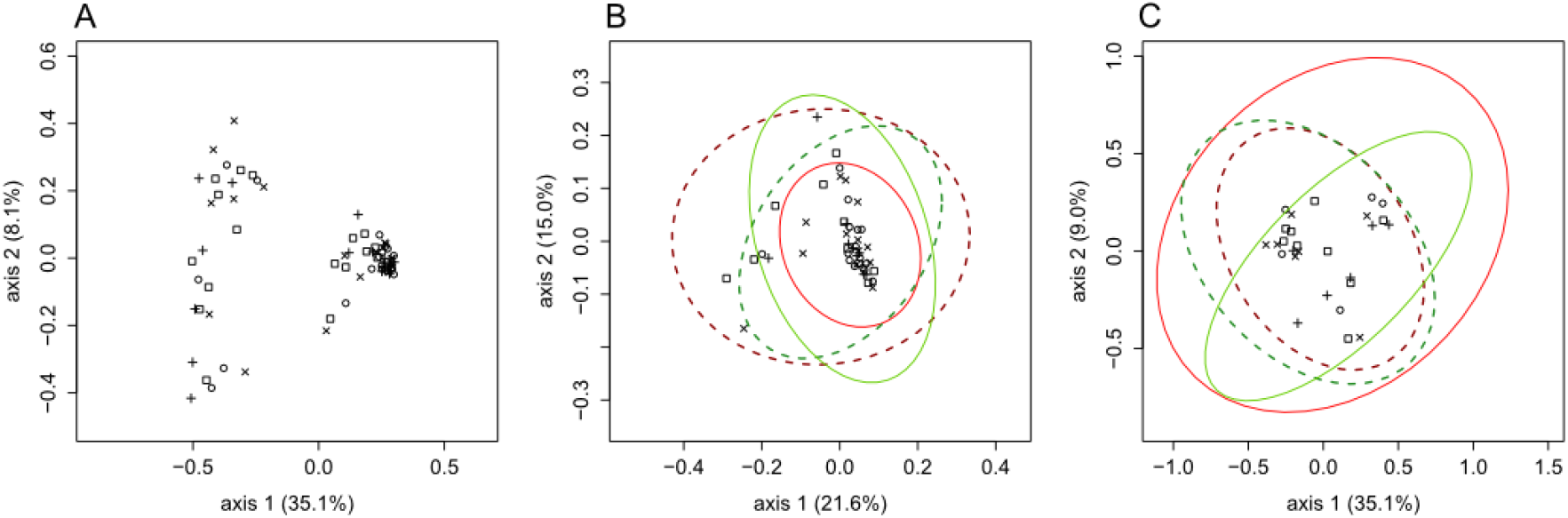
Principal coordinate analyses (PCoAs) between the microbiota of the four captive whitefish groups. A: comparison between the four captive whitefish groups intestinal microbiota. B: comparison between the four whitefish groups intestinal microbiota in the mixed diet group. C: comparison between the four whitefish groups intestinal microbiota in the *Artemia* diet group. Ellipses of 95% confidence were done with dataEllips using R car package. Each whitefish species is represented by different symbols: dwarf (D♀×D♂) and normal (N♀×N♂) are represented by circle a cross respectively and their ellipses are represented by continuous lines. The hybrid F1 N♀×D♂ and hybrid F1 D♀×N♂ are represented by the symbol × and □ respectively and their ellipses are represented by dotted line. Dwarf and hybrid F1 D♀×N♂ are represented in red whereas normal and hybrid F1 N♀×D♂ are represented in green.

We observed between eight and 42 bacterial genera specific to a given whitefish group in both diet groups (Fig. 6). We observed 21 dwarf-specific and 27 normal-specific bacterial genera respectively whereas the comparison between hybrids F1 D♀N♂ and F1 N♀D♂ revealed 41 and 16 specific bacterial genera respectively. Finally, 135 specific bacteria genera to the mixed diet group *versus* 62 to the *Artemia* diet group was also observed (see Table S4 for details).

**Figure 6.**
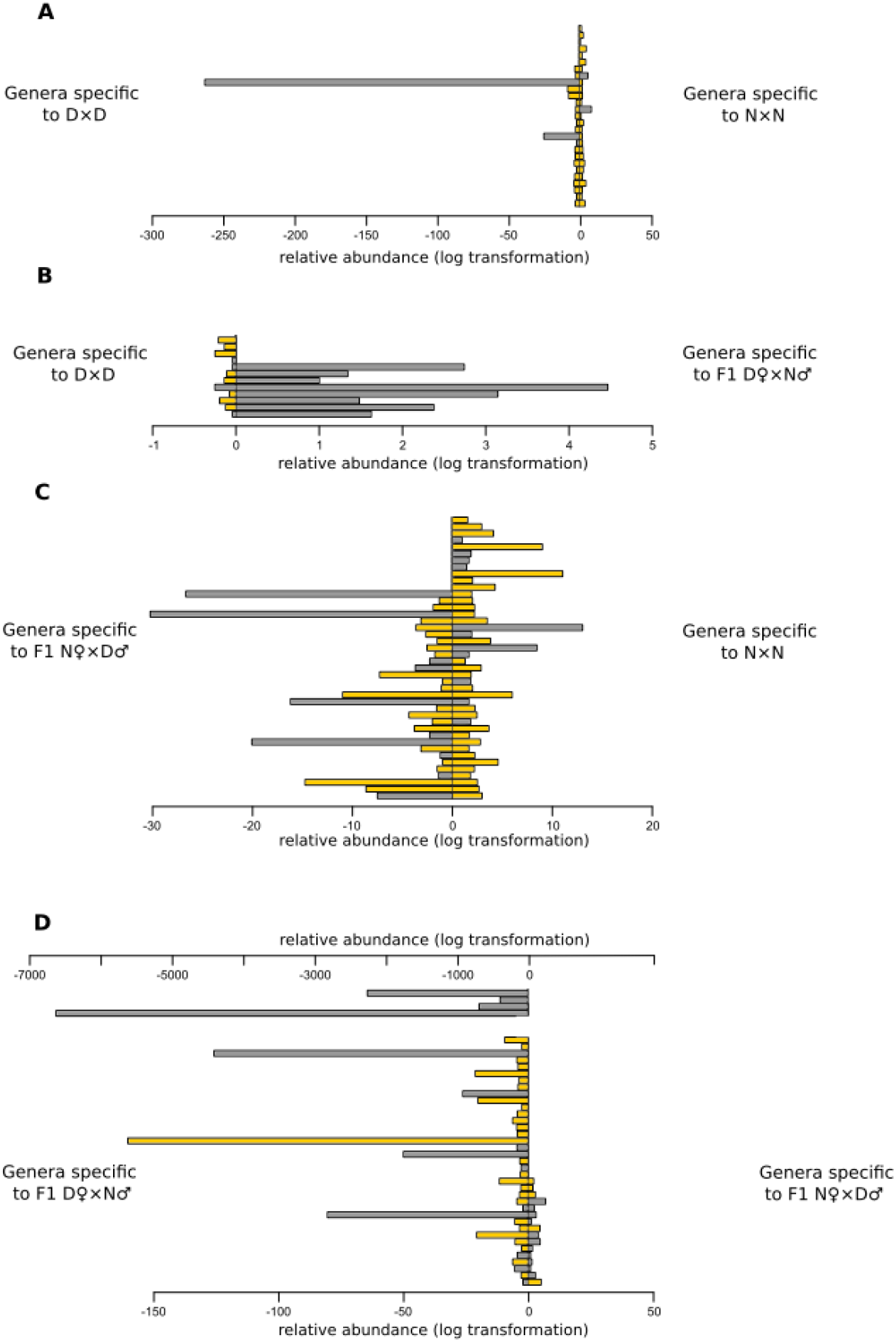
Metastats results for dwarf, normal and hybrid captive whitefish. Four side-by-side comparisons were performed with dwarf (D♀ × D♂), normal (N① × N♂), hybrid F1 N♀ × D♂ and hybrid F1 D♀ × N♂. The comparisons are log-transformed and each genus specific to a given whitefish group is represented by a bar plot. Mixed diet and *Artemia* diet groups are represented by yellow and grey bars, respectively.

## Discussion

In this study, we compared the transient intestinal microbiota among five wild sympatric species pairs of Lake Whitefish as well as captive dwarf, normal and hybrids whitefish reared in identical controlled conditions. More specifically, we tested for (i) differences in intestinal microbiota between dwarf and normal wild whitefish from the same lake; (ii) the occurrence of parallelism among five wild sympatric species pairs, and (iii) differences in intestinal microbiota among dwarf, normal and reciprocal hybrid whitefish which were reared together in identical controlled conditions.

### The intestinal microbiota of captive versus wild whitefish

Although an important part of bacteria which colonizes fish intestine may represent a random sampling from water and food, the occurrence of intestinal microbiota cores have been increasingly documented [58]. The intestinal microbiota cores represent OTUs or genera shared among closed host relatives. The comparison between wild and captive whitefish highlighted six genera shared by at least 80% of all samples. Interestingly, our intestinal core microbiota data represented 20% of shared sequences which is higher than the intestinal microbiota core reported for cichlid species (13-15%) [39]. These shared genera could be horizontally transmitted and/or selected as a common set of bacteria [39, 59]. Although the captive whitefish were hatched in captivity, their parents were of wild origin. Therefore, the conservation of certain genera by many captive whitefish might corroborate the microbiota vertical transmission in fish. Finally, it is noteworthy that we found many bacteria of unknown taxonomy (see Figure 2), and much more so in wild whitefish than in captive whitefish. This, along with previous studies, emphasizes that a considerable number of bacteria is waiting to be discovered in natural freshwater ecosystems.

### No clear pattern of parallel evolution in transient intestinal microbiota between dwarf and normal whitefish in the wild

Parallelism is the evolution of similar traits in independent populations [60] and has been well documented in several sympatric species from temperate and sub-arctic lakes throughout the north hemisphere [22, 61, 62]. Phenotypic parallelism between dwarf and normal whitefish has previously been documented for morphological, physiological, behavioral, and ecological traits [23, 25, 26, 28-31, 33, 63]. Given the difference in trophic and ecologic niches occupied by both species [25, 64], we predicted that some level of parallelism in transient intestinal microbiota would be observed between dwarf and normal whitefish species pairs. The dwarf whitefish is a limnetic fish feeding on zooplankton whereas the normal whitefish is a benthic fish feeding on zoobenthos and molluscs [24, 65]. Therefore, we expected that a different diet should bring the dwarf and normal whitefish of a given sympatric pair in contact with different bacterial communities, leading to a distinct transient intestinal microbiota in a similar manner in the different lakes. Moreover, differentiation of microbiota composition correlated with diet was previously observed [66-70]. In fact, the use of novel diet elements frequently produced a change in the microbiota composition by increasing or decreasing different bacterial strain according to their metabolic potential [11]. This is also supported by the microbiota composition differentiation of the two diet groups observed in captivity in this study. However, despite a global effect of species host on microbiota, our results revealed no clear pattern of parallelism among lakes. Indeed, difference between dwarf and normal whitefish microbiota composition was observed in three of the five lakes only whereas no difference was observed in the other two lakes. Together, this suggested that the environment has a more pronounced effect than the species host on the transient intestinal microbiota of dwarf and normal whitefish. These results are in line with those obtained in a previous study in the same system but investigating kidney microbiota [34]. Results of that study revealed that differences in bacteria composition between dwarf and normal whitefish were not parallel among lakes. However, unlike this study and in accordance with the higher diversity of prey types, normal whitefish kidney tissue consistently had a more diverse bacterial community and this pattern was parallel among lakes. Together, these results on whitefish microbiota add to building evidences from previous studies on this system that the adaptive divergence of dwarf and normal whitefish has been driven by both parallel and nonparallel ecological conditions across lakes. Moreover, the water bacterial community of the same studied lakes was investigated previously and we found that each lake is characterized by a specific water bacterial community [35]. This may reflect the differences in both biotic and abiotic factors among these lakes [25, 64]. More specifically, Cliff, Webster and Indian lakes are characterized by a greater oxygen depletion and a lower zooplankton densitiy whereas East and Témiscouata lakes are characterized by more favorable environmental conditions with a more important density of zooplankton and well-oxygenated water [64]. Therefore, the variation of water bacterial community along with the biotic and abiotic factors could underlie the more important lake effect than species host effect observed in the transient intestinal microbiota. Nevertheless, highly distinct bacterial composition between the water bacterial community and the whitefish transient intestinal microbiota was observed among lakes. The water bacterial community was dominated by *Proteobacteria*, *Actinobacteria* and *Bacteroidetes* whereas the whitefish transient intestinal microbiota was dominated by *Firmicutes* and *Proteobacteria* [35]. Therefore, whitefish transient intestinal microbiota was not directly reflective of its local environment, raising the hypothesis of a selective effect on microbiota induced by host physiology, immunity and genetic background [12, 18]. For instance, some transient bacteria might contribute to digestion of host diet [40] and such bacteria may impact on the transient intestinal microbiota composition by increasing their abundance [11].

### Comparison of transient and adherent intestinal microbiota in wild whitefish and the host effect

The most prevalent phyla in wild whitefish transient microbiota are *Acidobacteria, Actinobacteria, Bacteroidetes, Chlamydiae, Chloroflexi, Firmicutes, Fusobacteria, Planctomycetes, Proteobacteria, Terenicutes, TM7* and *Verrucomicrobia,* which have also been reported in previous studies of freshwater fishes [71-76]. In a previous study on adherent intestinal microbiota, (that is adherent to the intestinal mucosa) performed on the same individuals, we found that while the major phyla being represented were similar, the abundance of some of them was different [35]. For example, the five first phyla for the adherent microbiota were *Proteobacteria* (39.8%), *Firmicutes* (19%), *Actinobacteria* (5.1%), *OD1* (3.8%) and *Bacteroidetes* (2.8%) whereas the five first phyla for the transient microbiota were *Firmicutes* (38.2%), *Proteobacteria* (29.5%), *Verrucomicrobia* (4.4%), *Planctomycetes* (4.1%) and *Actinobacteria* (3.7%). Moreover, the number of genera and the number of OTUs were much more important in the transient microbiota (611 genera and 94,883 OTUs) than the adherent microbiota (421 genera and 10,324 OTUs). Most of the adherent bacterial taxa living on the intestinal mucosa are not randomly acquired from the environment [77], but are rather retained by different characteristics of the host [13]. It was also reported that there is an important host effect in both dwarf and normal whitefish, which stabilizes the number of bacterial genera living in the intestinal mucosa [35]. Thus, the comparison between whitefish transient and adherent microbiota supports the view that the whitefish host have a substantial selective effect on its intestinal microbiota. For instance, dwarf and normal whitefish in Cliff and East lakes show a distinct intestinal microbiota for both the adherent and the transient bacteria. It is also noteworthy that the adherent, but not the transient intestinal microbiota differed between species in Témiscouata Lake whereas the exact opposite was observed in Indian Lake. In Témiscouata Lake, this difference in adherent microbiota between species suggested a host-species effect leading to differential abundance of the same bacterial taxa. In contrast, results in Indian Lake suggest that host species have no clear effect on microbiota divergence and that the difference in transient microbiota is likely caused by the trophic niches occupied by each species. Altogether, these observations suggest that the direction and intensity of factors determining the composition of intestinal microbiota may differ between the host and the microbiota of a given holobiont system, as previously reported [7]. Here, three distinct host-microbiota interactions may have evolved independently in postglacial time: (i) divergence of intestinal microbiota influenced by the host and the environment (Cliff and Est lakes), (ii) divergence of the intestinal microbiota mostly influenced by the host (Témiscouata Lake), (iii) divergence of intestinal microbiota mostly influenced by the environment (Indian Lake). Finally, given the pronounced difference that may exist between transient and adherent microbiota, our results suggest that adherent microbiota is a more reliable choice to study the effect of host species on microbiota than the analysis of transient microbiota.

### Modest but significant host effect on the transient intestinal microbiota in controlled conditions

An unplanned variation in our experimental set up occurred during the captive rearing of the whitefish pair species and the reciprocal hybrids during seven months, which led to the unexpected observation of a diet preference which split the whitefish into two groups independently of the parental or hybrid origin or the tanks where fish were. The use of two types of food, *Artemia* and dry pellets, are usually recommended for optimizing growth and survival of juvenile whitefish in captivity [43, 44]. However, while 47 whitefish opted to feed on both types of diet, 27 chose to feed only on *Artemia.* This allowed us to assess the impact of different diets in an otherwise identical controlled environment, which revealed that diet had the most profound impact on the community composition of transient intestinal microbiota in a controlled environment.

Nevertheless, we did observe a significant, albeit modest effect of whitefish genetic hosts on the transient intestinal microbiota. In principle, in a controlled environment, there should be no environmental effect on the microbiota composition and consequently, variation in microbiota should only depend on the host effect which integrated the influence of the host physiology, immunity, as well as genetic background. Here, while the PCoA analysis only revealed a slight pattern of differentiation between both parental species and their reciprocal hybrids, the PERMANOVA test revealed a statistically significant effect of the host genetic background on the taxonomic composition of the transient microbiota. This was accompanied by a significant variation in bacterial abundance at the phylum level, especially within the diet group A (feeding on both *Artemia* and dry pellets). Finally, numerous genera that were specific to one whitefish species or the hybrids were observed in both diet groups. These results suggest an effect of hybridization on the transient intestinal microbiota. This hybridization effect could be explain by the genetic incompatibilities model of Bateson, Dobzhansky and Muller (BDM) previously highlighted in the Lake Whitefish system [63, 78, 79]. To our knowledge, only one study compared the intestinal microbiota among closely related fish populations in controlled conditions [38] and none compared parental and hybrid progeny. Specifically, distinct intestinal microbiota between two ecotypes of the Trinidadian Guppy (*Poecilia reticulata*) suggested a pronounced effect of the genetic background [38]. However, these results should be interpreted cautiously since fish used for this experiment were adults that were born in the wild and kept in tanks for 10 weeks only. Consequently, the difference could reflect a carry-over effect from the natural conditions whereas in our case, fish were born in captivity.

## Conclusions

In summary, we observed complex patterns of variation in the transient intestinal microbiota of sympatric dwarf and normal whitefish in their natural environment, with little evidence of parallelism among different populations of a same species. Local environment, more than species explained the observed patterns of variation. Nevertheless, a possible host selective effect was revealed by the pronounced contrast in the bacterial composition observed in water and intestinal samples. Moreover, differences in taxonomic composition between sympatric species were observed for three of the five lakes, suggesting specific but not parallel microbiota-host association in these lakes. Finally, a possible host effect on its transient intestinal microbiota was revealed by a modest, albeit significant genetic host effect (parental forms and reciprocal hybrids) when reared in controlled conditions. However, a more prevalent effect of environmental factor was highlighted by the pronounced difference observed among two diet groups. Overall, our results support the view that the transient intestinal fish microbiota is the result of complex interactions between the host’s genetic background and environmental conditions such as trophic resources. The fact that we observed stronger environmental effect on the microbiota among five sympatric whitefish pairs illustrates that drawing generalization regarding host-microbiota association for a given species may be difficult, and in fact inappropriate.

## Acknowledgements

We thank G. Côté, A. Dalziel, A-M. Dion-Côté, S. Higgins and J-C Therrien for fieldwork and technical assistance for crossing and rearing of captive whitefish at the LARSA. We are grateful to C. Hernandez-Chàvez for laboratory advice and support, B. Boyle for his help with the Illumina MiSeq sequencing.

## Supplementary Data

### Experimental crosses of captive whitefish

Whitefish eggs used for this study were incubated at the Laboratoire de Recherche en Sciences Aquatiques (LARSA, Université Laval, Québec, Canada). The dwarf species came from Témiscouata Lake (47°40’04”N, 68°49’03”W) which is from the Acadian glacial lineage origin whereas the normal species came from Aylmer Lake (45°54”N, 71°20”W) corresponding to the Atlantic glacial lineage [23]. Backcross F1-Hybrids were obtained by crossing a F1 hybrids laboratory strain and wild whitefish parents. More precisely, F1 hybrid (F1 D ♀ *N♂) were produced in crossing three wild dwarf females and two laboratory strain normal males by artificial fertilization. Same processes was used to produced F1 hybrid (F1 N ♀ *D♂) with crossing five laboratory strain normal females and twelve wild dwarf males (see figure 1 [80]). The dwarf and normal whitefish crosses were also created by artificial fertilization with sperm and eggs were collected in the field and transported to the LARSA. No treatments, such as antibiotics or malachite green were delivered to the eggs.

### Whitefish host: DNA extraction, amplification and genetic identification of captive whitefish lineages

A fin clip was collected from all fish and DNA was extracted using a salt extraction method [81] with slight modifications [82]. Mitochondrial (mtDNA) and nuclear DNA were used to identify the whitefish dwarf and normal, and their hybrids (F1 hybrid D♀N♂ and F1 hybrid N♀D♂). First, an analysis of mtDNA restriction fragment length polymorphism (RFLP) was performed as described in Dalziel et al. since pure dwarf and normal species possess distinct mitochondrial DNA haplotypes [29, 83]. In brief, after the amplification of the cytochrome b by PCR, the amplified products were digested with SnaBI which cuts the amplified cytochrome b of the normal whitefish haplotype but not of the dwarf. Second, 12 nuclear microsatellite loci were genotyped on all juvenile whitefish and their known parents to differentiate them at the nuclear DNA level and details about primer sequences and PCR protocols are presented in Rico et al.. Three different PCRs were performed for this whitefish microsatellite markers analysis [84]. Firstly, the multiplex PCR A was performed with 2 μl (≈20 ng) of whitefish DNA, 5 μL Qiagen^®^ multiplex reaction buffer, forward and reverse primers at different concentrations: 0.3 μm of Cocl32, Cocl lav41, Cocl Lav8 and 0.35 μm of Cocl Lav224; purified water adjusted the final volume at 10 μl. Multiplex PCR program was: 15 min at 94°C, and then 35 cycles of 30 sec at 94°C, 3 min at 58°C, 1 min at 72°C and 30 min at 60°C. Secondly, the multiplex PCR B were performed with 2 μl (≈20 ng) of whitefish DNA, 5 μL Qiagen^®^ multiplex reaction buffer and forward and reverse primers at different concentration: 6 μm of Cocl15 et Cisco200 and 0.25 μm of Cocl 33; purified water adjusted the final volume at 10 μl. Multiplex PCR program was: 15 min at 94°C, and then 35 cycles of 30 sec at 94°C, 3 min at 60°C, 1 min at 72°C and 30 min at 60°C. Thirdly, the Simplex PCRs were performed with 2 μl (≈20 ng) whitefish DNA, 0.2 μl GoTaq^®^ DNA polymerase (PROMEGA), 0.5 μl of each forward and reverse markers (0.5 μm) (Osmo5, Cocl34, Cocl36, Bwf F-1 and Cocl Lav22) 2 μl of 5X Colorless GoTaq^®^, 0.6 μl of MgCl2 (0.5 mM), 0.8 μl dNTPs (200 μm) and purified water adjusted the final volume at 10 μl. Simplex PCR program was: 2 min at 94°C, and then 35 cycles of 30 sec at 94°C, 3 min at 58°C (Osmo5, Cocl36, Bwf F-1, Cocl Lav22) or 64°C (Cocl34), 1 min at 72°C and 30 min at 60°C. Amplified loci were migrated via electrophoresis using an ABI 3130xl capillary DNA sequencer (Applied Biosystems Inc.) with a molecular size standard (GeneScan-500 LIZ, Applied Biosystems). Genotypes were scored using Genemapper 4.0 (Applied Biosystems Inc). A combination of three software, STRUCTURE v2.3.4, GENECLASS2 v2.0 and PAPA v2.0 was used to reassign each studied fish to its group of origin [85-87]. STRUCTURE was performed assuming an admixture model without priors with a burn-in period of 50 000 followed and 100,000 Markov Chain Monte Carlo (MCMC) steps. GENECLASS2 was conducted using the simulation test of [88] based on 100,000 simulated individuals. Finally, PAPA was performed for the parental allocation procedure with a uniform error model (error sum = 0.02).

**Table S1.** Steps used to reduce sequencing and PCR errors. We followed the step recommended by MOTHUR in the MiSeq SOP protocol.

| Main Step of filtration | Number of<br>microbiota<br>wild<br>whitefish<br>sequences | Number of<br>microbiota<br>captive<br>whitefish<br>sequences | Number of<br>microbiota<br>captive and<br>wild<br>whitefish<br>sequences |
| --- | --- | --- | --- |
| After contigs construction | 4765128 | 2498271 | 8299965 |
| Remove sequences with ambiguous bases and lengths more than 450 bp | 1774729 | 885737 | 2660466 |
| Aligning paired ends (maximum two mismatches) and remove sequences with homopolymers of more than eight bp and with lengths less than 400 bp | 1737037 | 872921 | 2609958 |
| Remove chimeric sequences | 1729598 | 868868 | 2598519 |
| Remove sequences from chloroplasts, mitochondria and nonbacterial | 1855778 | 845370 | 2498271 |
| Number of final sequences | 1855778 | 845370 | 2498271 |
| Number of OTUs | 94883 | 85363 | 189683 |
| Number of genus | 611 | 433 | 710 |
| Good's Coverage | 95.06% | 88.22% | 91.86% |

**Table S2.** Matrix of bacterial abundance and Good’s coverage per captive whitefish sample. DD: dwarf whitefish, NN: normal whitefish, DH: hybrid F1 D♀*N♂, NH: F1 N♀*D♂. The diet group A is composed of *Artemia* and dry food; B is composed of *Artemia*.

| Group | Whitefish species | Body mass (gram) | Diet Group | Number of sequences | Good's coverage |
| --- | --- | --- | --- | --- | --- |
| 04-B2F | NH | 62.50 | A | 9554 | 83.63 |
| 04-B3F | NH | 68.45 | A | 4047 | 78.95 |
| 05-B1F | NN | 54.11 | A | 7950 | 87.38 |
| 05-B2F | DD | 48.75 | A | 4272 | 77.76 |
| 05-B3F | NH | 49.62 | A | 10535 | 84.72 |
| 06-B1F | DH | 20.27 | A | 10626 | 86.82 |
| 06-B2F | NH | 53.55 | A | 9598 | 85.02 |
| 07-B1F | DD | 70.76 | A | 10220 | 83.05 |
| 07-B3F | NN | 66.45 | A | 12571 | 88.09 |
| 08-B3F | NH | 38.24 | A | 4981 | 84.08 |
| 09-B1F | DD | 35.43 | A | 8830 | 86.87 |
| 09-B3F | NN | 78.10 | A | 9552 | 78.43 |
| 10-B1F | DD | 66.88 | A | 8965 | 82.96 |
| 10-B3F | DD | 40.19 | A | 5368 | 80.61 |
| 11-B1F | DD | 70.33 | A | 9494 | 83.98 |
| 11-B3F | DD | 15.23 | A | 4764 | 76.20 |
| 12-B1F | NH | 41.27 | A | 2616 | 81.04 |
| 12-B2F | DH | 12.24 | A | 11443 | 81.56 |
| 12-B3F | NH | 96.84 | A | 8438 | 84.55 |
| 13-B2F | DH | 15.65 | A | 8778 | 84.43 |
| 14-B1F | NH | 45.23 | A | 7496 | 80.31 |
| 14-B3F | DD | 61.29 | A | 8842 | 80.32 |
| 15-B1F | DH | 26.03 | A | 9462 | 86.81 |
| 15-B2F | NN | 97.68 | A | 9109 | 82.89 |
| 15-B3F | DH | 51.04 | A | 7660 | 79.13 |
| 16-B3F | NH | 70.71 | A | 7866 | 78.85 |
| 17-B1F | NN | 64.96 | A | 4621 | 78.84 |
| 17-B3F | DD | 86.65 | A | 7478 | 80.82 |
| 18-B2F | DD | 78.20 | A | 7387 | 81.18 |
| 19-B3F | NN | 83.91 | A | 14935 | 82.68 |
| 20-B1F | DD | 22.63 | A | 9012 | 80.42 |
| 21-B2F | NN | 89.88 | A | 9285 | 82.16 |
| 21-B3F | NN | 49.14 | A | 10663 | 83.77 |
| 22-B1F | NH | 46.79 | A | 10264 | 85.83 |
| 22-B2F | DH | 19.08 | A | 14523 | 82.39 |
| 23-B3F | DH | 59.35 | A | 5548 | 81.65 |
| 24-B1F | DH | 19.75 | A | 12012 | 83.70 |
| 24-B2F | DD | 62.52 | A | 3249 | 75.62 |
| 24-B3F | DD | 3.60 | A | 7171 | 80.83 |
| 25-B1F | NH | 63.67 | A | 10637 | 81.66 |
| 26-B2F | NN | 32.98 | A | 7388 | 83.05 |
| 27-B2F | NH | 57.57 | A | 11992 | 91.12 |
| 27-B3F | DD | 45.78 | A | 12970 | 88.66 |
| 29-B1F | NH | 55.69 | A | 16608 | 84.97 |
| 30-B2F | DD | 41.33 | A | 8130 | 83.89 |
| 33-B2F | DH | 2.99 | A | 7908 | 90.59 |
| 33-B3F | DH | 28.31 | A | 7948 | 81.35 |
| 02-B1F | NH | 58.59 | B | 7773 | 96.57 |
| 03-B3F | NH | 64.97 | B | 14847 | 99.51 |
| 06-B3F | DH | 3.59 | B | 9042 | 98.83 |
| 07-B2F | DD | 2.82 | B | 3941 | 97.64 |
| 08-B1F | DD | 6.08 | B | 16447 | 98.61 |
| 08-B2F | NH | 3.44 | B | 12825 | 99.27 |
| 09-B2F | NH | 5.80 | B | 24288 | 96.83 |
| 10-B2F | DH | 3.55 | B | 20371 | 99.36 |
| 13-B1F | DH | 2.93 | B | 7346 | 94.80 |
| 16-B1F | DH | 6.29 | B | 13921 | 98.99 |
| 18-B1F | DH | 1.57 | B | 10261 | 94.29 |
| 19-B2F | NN | 3.09 | B | 16329 | 98.51 |
| 20-B2F | NH | 57.16 | B | 13028 | 98.53 |
| 20-B3F | NH | 7.57 | B | 14615 | 98.47 |
| 21-B1F | NN | 2.85 | B | 6503 | 97.91 |
| 22-B3F | DD | 3.01 | B | 9433 | 99.14 |
| 23-B1F | DH | 2.41 | B | 11224 | 97.46 |
| 25-B2F | DH | 2.65 | B | 19484 | 99.12 |
| 25-B3F | DD | 70.32 | B | 15942 | 99.43 |
| 26-B1F | NN | 2.85 | B | 11586 | 99.07 |
| 26-B3F | NN | 3.55 | B | 14829 | 98.92 |
| 28-B3F | NN | 3.71 | B | 17491 | 99.40 |
| 29-B3F | DH | 2.71 | B | 7009 | 97.73 |
| 30-B3F | DH | 3.12 | B | 28573 | 99.30 |
| 31-B1F | DD | 7.26 | B | 17016 | 96.21 |
| 31-B3F | NH | 2.50 | B | 12031 | 99.13 |
| 32-B1F | NN | 4.77 | B | 13956 | 98.64 |

**Table S3.** Summary of ANOVA test statistics on microbiota alpha diversity (inverse Simpson index). All lakes refer to the analysis of effect of host species (dwarf and normal), lake (Cliff, East, Indian, Témiscouata and Webster). Second, the effect of host species is treated for each lake separately. Third, CAPTIVE refers to the analysis of effect of host group (dwarf, normal, hybrids D♀N♂ and N♀D♂), diet (*Artemia* only and *Artemia* with dry food) on all captive fish. Fourth, effect of captivity (wild and captive), dwarf only and normal only. F-Value is the value of the F statistic.

| Fish group | Source of variation | Degrees of freedom | F value | P value |
| --- | --- | --- | --- | --- |
| WILD |  |  |  |  |
| All lakes | Species | 1 | 0.439 | 0.510 |
|  | Lakes | 4 | 2.304 | 0.064 |
|  | Lakes:Species | 4 | 1.152 | 0.337 |
| Cliff | Species | 1 | 0.109 | 0.744 |
| Est | Species | 1 | 0.025 | 0.876 |
| Indian | Species | 1 | 2.026 | 0.169 |
| Témiscouata | Species | 1 | 1.557 | 0.225 |
| Webster | Species | 1 | 0.824 | 0.380 |
| CAPTIVE |  |  |  |  |
|  | Group | 3 | 0.599 | 0.620 |
|  | Diet | 1 | 40.471 | 0.001 |
|  | Species:Diet | 3 | 1.930 | 0.134 |
| BOTH |  |  |  |  |
| All fish | Captivity | 1 | 8.915 | 0.003 |
| Dwarf | Captivity | 1 | 2.044 | 0.157 |
| Normal | Captivity | 1 | 2.040 | 0.157 |

**Table S4:**
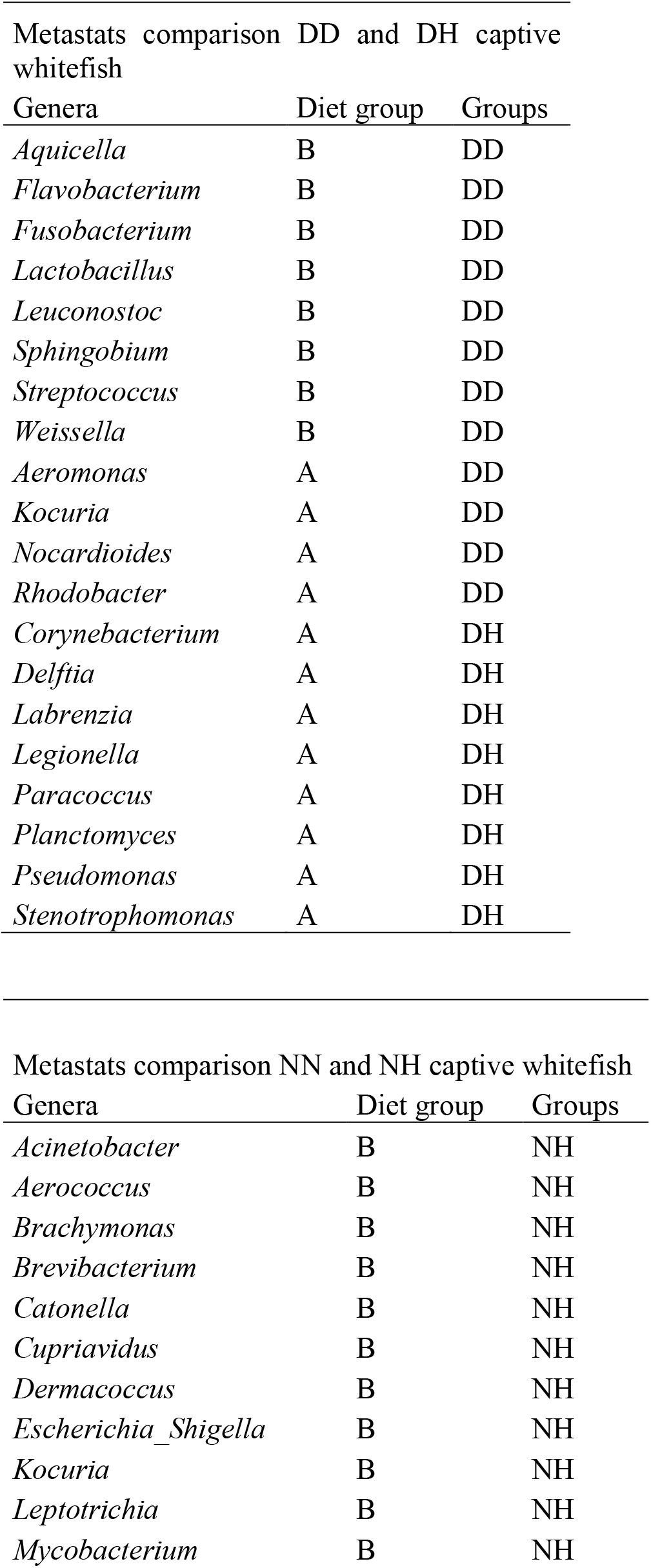
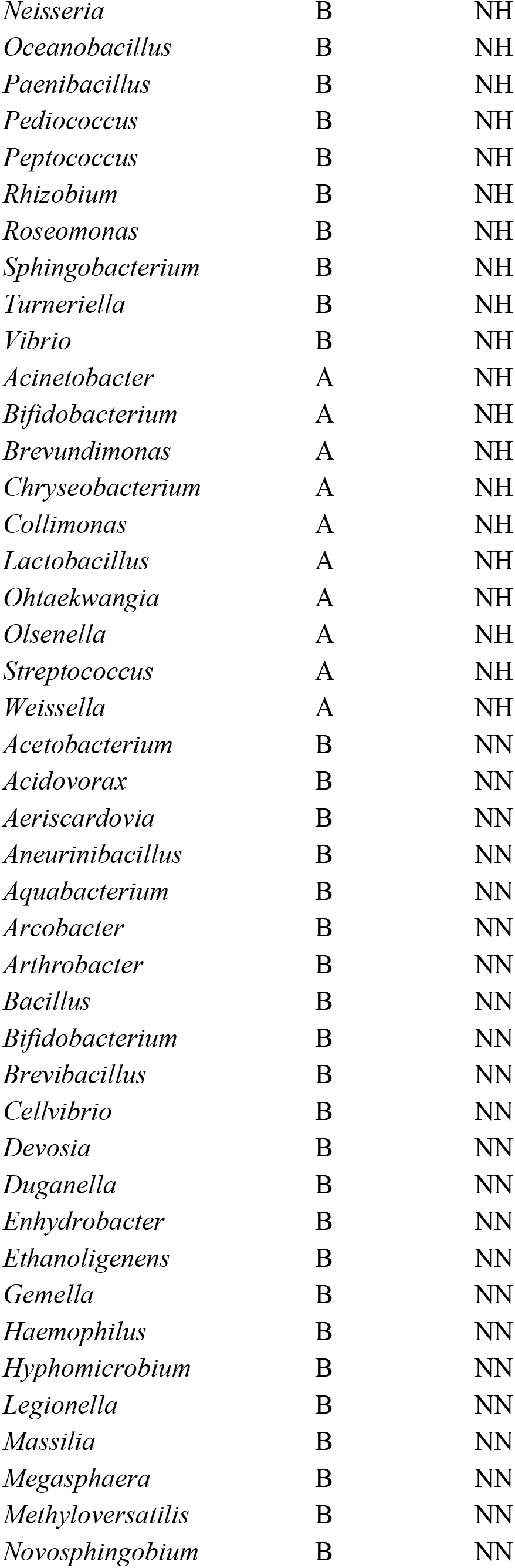
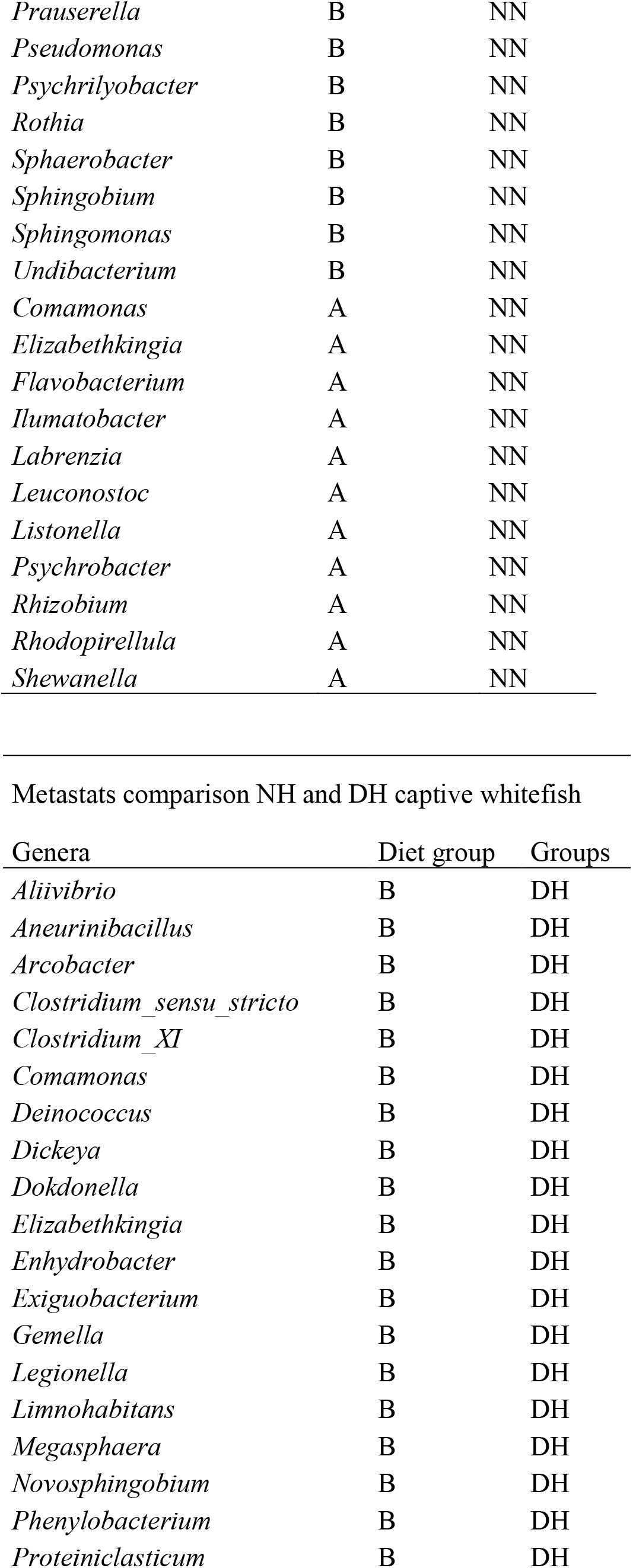
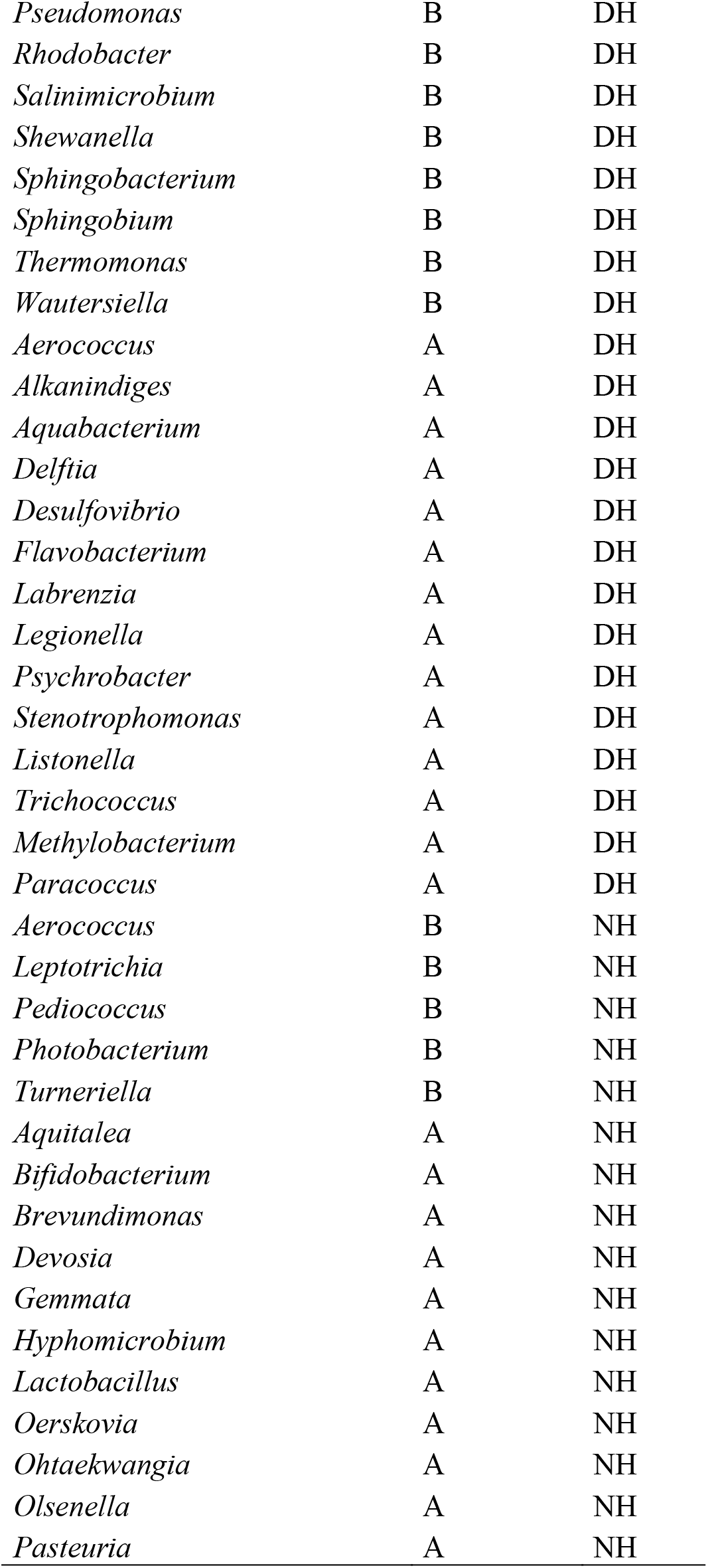
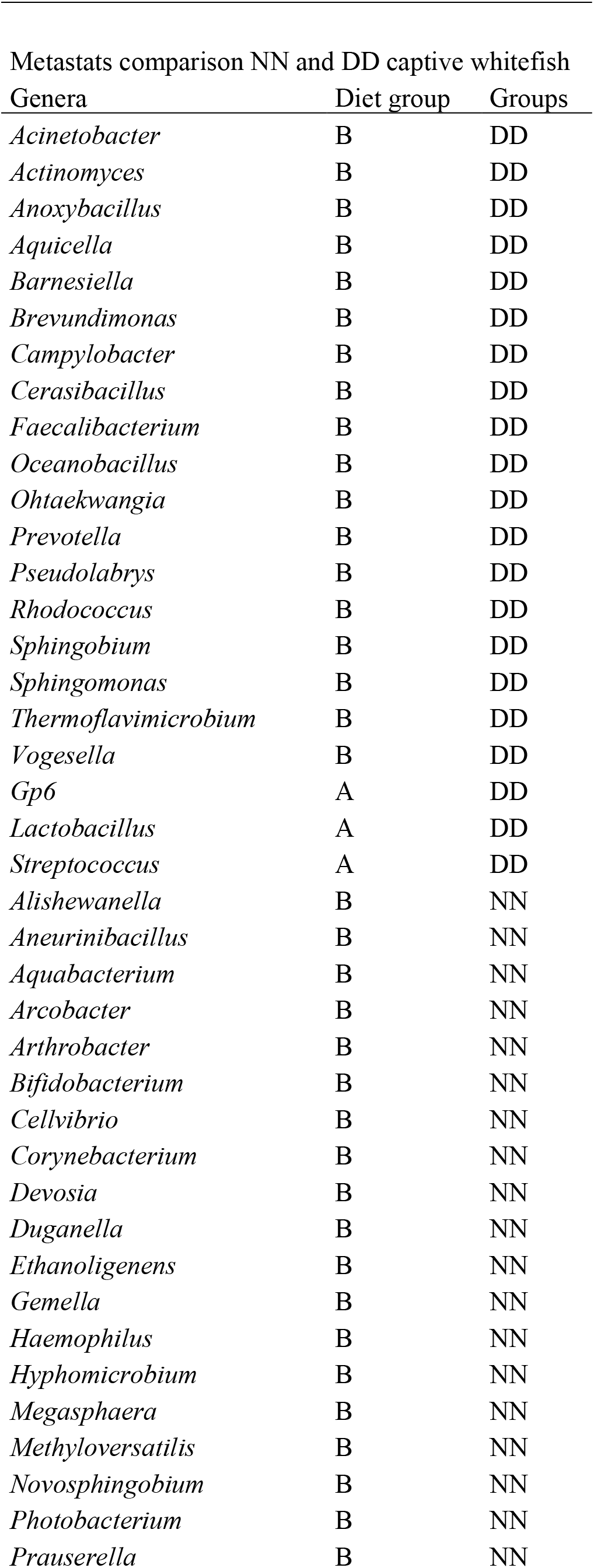
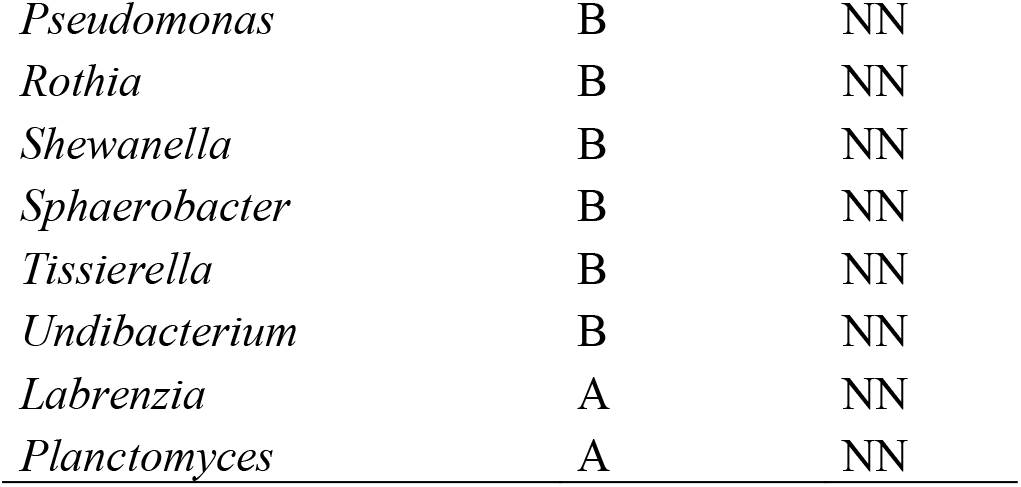
Four Metastats tables with details of one-species-specific genera. DD: dwarf whitefish, NN: normal whitefish, DH: hybrid F1 D♀*N♂, NH: F1 N♀*D♂. The diet group A is composed of *Artemia* and dry food; B is composed of *Artemia.*

**Figure S1.**
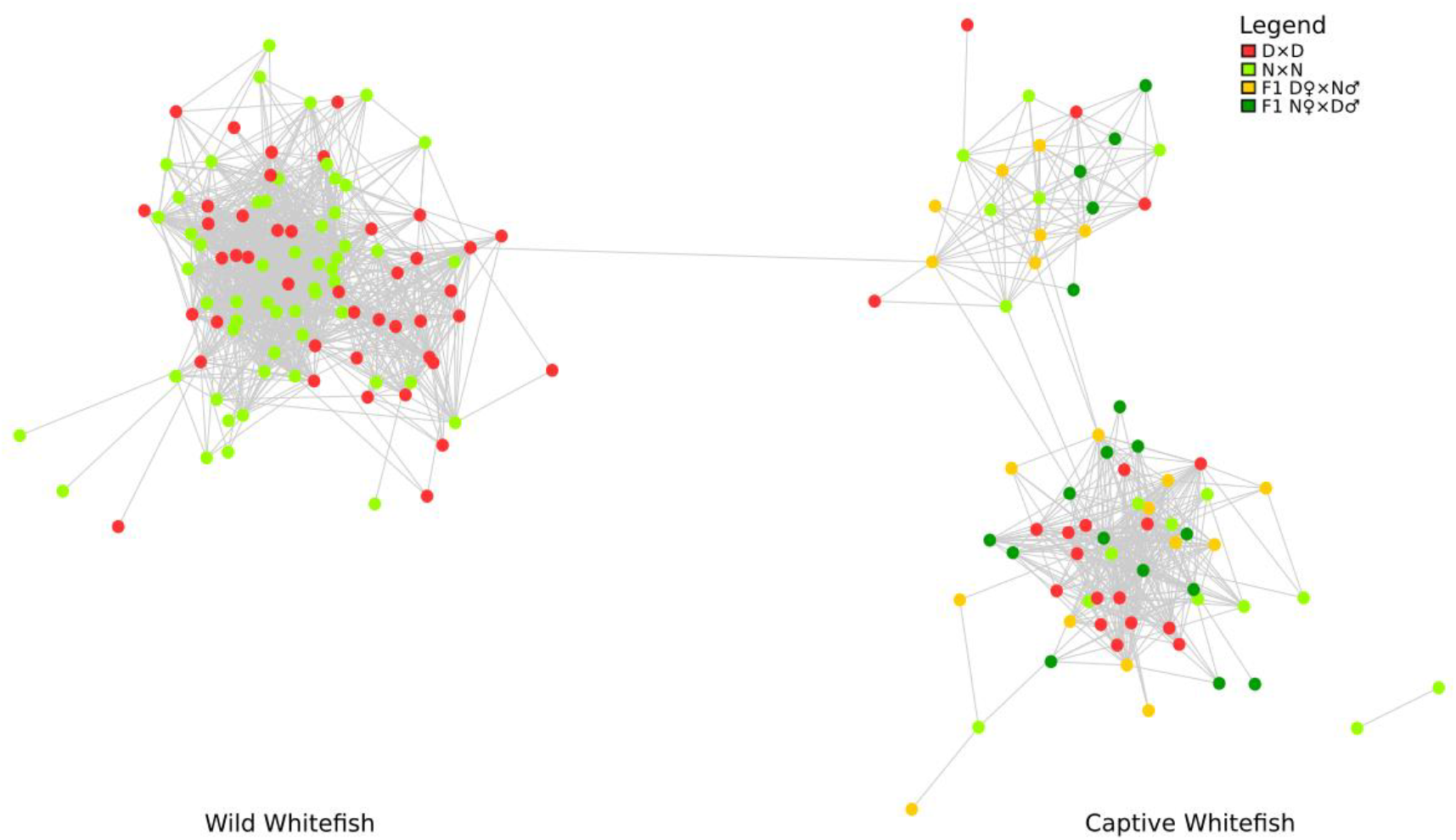
Network analysis of intestinal microbiota of dwarf and normal wild whitefish and intestinal microbiota of dwarf, normal and hybrids captive whitefish. The nodes represent a dwarf or a normal or a hybrid whitefish microbiota. More precisely, DD: dwarf whitefish, NN: normal whitefish, DH: hybrid F1 D①*N♂, NH: F1 N①*D♂. The connecting lines between two samples represent their correlation and is highlighting by a Spearman index.

